# Evaluating a Primary Healthcare Centre’s Preparedness for Disasters Using the Hospital Safety Index: Lessons Learned from the 2014 Floods in Obrenovac, Serbia

**DOI:** 10.1101/503557

**Authors:** Zoran Lapčević, Stefan Mandić-Rajčević, Milan Lepić, Mića Jovanović

**Author notes:** **Corresponding Author:** Stefan Mandić-Rajčević, M.D., Ph.D., Innovation Center of Faculty of Technology and Metallurgy, University of Belgrade, Karnegijeva 4, 11000 Belgrade, Serbia.

## Abstract

Various organizations have endeavored to develop assessment methods for the identification and management of weaknesses in hospital disaster preparedness. Although the largest number of patients receive their regular care at the primary level, there is no internationally validated tool for the rapid safety assessment of primary health care centers (PHC). Flooding accounts for almost 50% of all disasters related to weather, and climate models consider these events as highly probable in the future. In May 2014, heavy rain caused floods affecting around 1.6 million people in Serbia, leaving the municipality of Obrenovac most severely impacted. This paper aims at assessing the safety of PHC Obrenovac using the Hospital Safety Index (HSI), evaluating the usefulness of HSI for safety assessment of PHCs, and drawing lessons from the 2014 floods. PHC Obrenovac had an overall safety index of 0.82, with structural, nonstructural safety, and disaster management indices of 0.95, 0.74, and 0.75, respectively, implying it is likely to function in disasters. A detailed analysis of individual HSI items underlined the necessary improvements in the field of emergency power and water supply, telecommunication, and emergency medical supplies, which rendered the PHC nonfunctional during the 2014 floods. Most items were considered of same relevance for primary healthcare centers as for hospitals, excluding some items in the medical equipment, patient care, and support services. Fine-tuning the HSI to primary healthcare settings, officially translating it into different languages, facilitating scoring and analysis could result in a valid safety evaluation tool of primary healthcare facilities.

**Highlights:**

- The Hospital Safety Index can be modified and used for primary health care centers
- The evaluated primary health care Centre is likely to function in disasters
- HSI identified flaws which would disable the PHCs functioning in case of floods
- Most improvements are necessary in the emergency power and water supply categories

## 1. Introduction

The Hyogo Framework for Action calls to *“Integrate disaster risk reduction into the health sector… and implement mitigation measures to reinforce existing health facilities, particularly those providing primary health care”* [1]. As one of the priorities at the national and local levels, the Sendai Framework for Disaster Risk Reduction 2015-2030 recommends *“To enhance the resilience of national health systems, by integrating disaster risk management into primary secondary and tertiary health care, especially at the local level…”* [2]. It is the job of local governments to provide essential services to their citizens and communities, such as health care, which need to be resilient to disasters [3]. Although the largest number of patients receive their regular care at the primary level, there is no internationally validated tool for the rapid safety assessment of primary health care centers (PHC) [4–6].

Different scientists and organizations have put an effort to develop assessment methods to facilitate the identification and management of weaknesses in hospital disaster preparedness. Higgins et al. (2004) assessed preparedness of hospitals in Kentucky (USA) using an instrument based on the Mass Casualty Disaster Plan Checklist [7]. Adini et al. (2006) analyzed various models for assessing the emergency preparedness of hospitals in mass-casualty incidents [8]. A comparison of an on-site survey, directly observed drill performance, and video analysiss of teamwork was done in 6 Los Angeles County hospitals by Kaji et al (2008) [9]. Lazar et al. (2009) endorse the use of measurable, evidence-base benchmarks and objective standards in hospital emergency management [10]. Top et al. (2010) examined the disaster plans of hospitals throughout Turkey using this method to estimate the preparedness for possible disasters [11]. World Health Organization (WHO) has developed the Hospital Safety Index (HSI), which is a validated, international, multi-risk assessment tool which allows for standardized comparisons of hospital safety levels [12]. The HSI has been used to assess hospital safety around the world, and studies evaluating up to several hundred hospitals have been published [13–16]. In Italy, Aiello et al. (2012) developed a simplified methodology based on the HSI to map the seismic risk for hospital buildings taking into account the specific national features, while Miniati and Iasio (2014) proposed a methodology which considered the complexity of the hospital system while leveraging the rapid assessment provided by the WHO evaluation forms, and applied it to 5 most important hospitals in the Province of Florence in the scenarios of earthquakes and floods [17,18].

The preparation to care for populations with chronic health conditions during disasters has been identified as a key issue in disaster preparedness [19]. The high burden of chronic diseases such as hypertension and diabetes has emphasized the need to develop disaster planning for such populations [19–23]. In a recent literature review regarding primary health care and disasters, Redwood Campbell et al (2011) underline the difficulty of defining “primary health care”, with the most common definition including care provided by physicians and activities such as “gatekeeper” (access to secondary and tertiary care), immunizations, prescriptions, and provision of basic medical services. The authors emphasize a lack of literature focusing on DRR, preparedness and recovery concerning PHCs [24]. There is only one paper dealing with the risk assessment of a PHC’s service interruption during a disaster, providing a simplified and specific assessment procedure for the flood hazard in Sudan [25].

Floods have resulted in extensive mortality and morbidity throughout the world, and can be considered one of the most common natural disaster [26,27]. Between 1995 and 2004 there has been an average of 127 floods per year, rising to 171 between 2005 and 2014. This number accounted for almost 50% of all disasters related to weather [28]. It has been estimated that flooding has taken more than 200,000 lives and affected almost 3 billion people worldwide [5]. Climate models consider high volume of rainfall and consequent flooding events highly probable in the future, thus underlining the importance of this kind of disaster [29,30]. Information on the influence of floods on the primary health care systems is lacking, and has not been well documented [5,31]. In the second half of May 2014, a low-pressure system named “Yvette” caused heavy rains to fall on Serbia. In one week, rainfall equivalent to 3 months of rain under normal conditions caused a rapid and substantial increase in water levels of main rivers in this area. It was estimated that the floods had affected around 1.6 million of people living in 38 municipalities/cities mostly located in central and western Serbia, among which the municipality of Obrenovac was most severely impacted [32].

This study aims at assessing the safety and disaster preparedness of the Obrenovac PHC using the HSI, evaluating the usefulness of HSI for the safety assessment of PHCs, and discussing these results taking into account this health care facility’s functioning during the 2014 floods.

## 2. Methods

### 2.1. The setting

***Figure 1*** shows the position of Serbia in Europe, as well as the position of Obrenovac in Serbia. Obrenovac is located around 30 km south-west of Belgrade (capital city of Serbia), near river Sava to the north. It is a suburban municipality of the city of Belgrade, with a total population of around 71,000, of which more than 24,000 live in the urban area. The primary health care Centre (PHC) Obrenovac, founded in 1952, covers 29 smaller towns and an area of 410 km^2^, with the most distant village 33 km away. It employs 337 healthcare workers, of which 113 are medical doctors.

**Figure 1.**
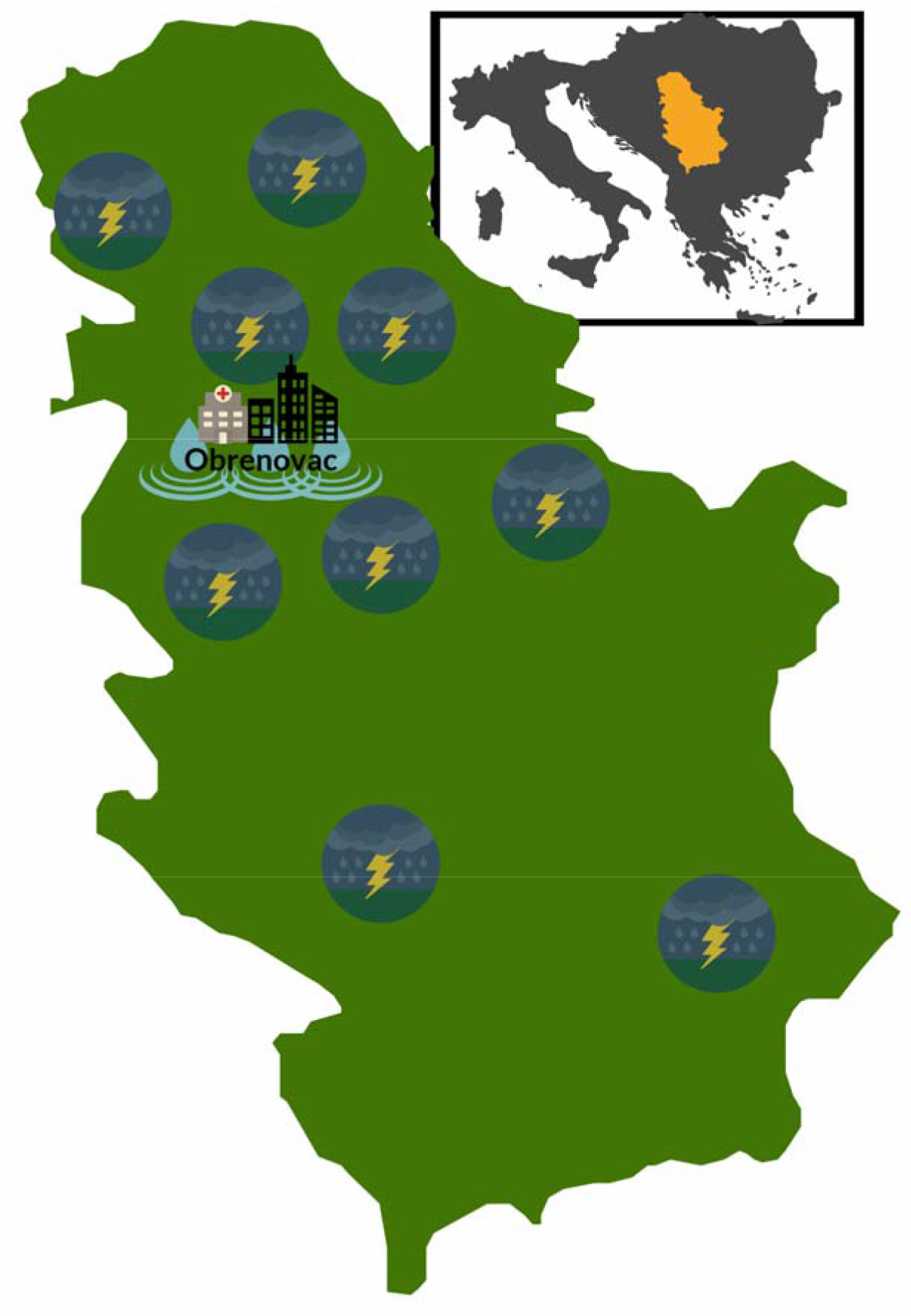
The position of Serbia in Europe and the position of Obrenovac in Serbia.

### 2.2. Hospital Safety Index

The Hospital Safety Index (HSI) is a tool for rapid, reliable and cost-effective diagnostic of the structural safety, non-structural safety and functional capacity of a hospital in 151 areas [12]. The 151 variable, each with three safety levels (***Low*** = “Unlikely to function”, ***Average*** = “Likely to function”, and ***High*** = “Highly likely to function”) are divided into four sections/modules: geographic location of the hospital, structural safety, nonstructural safety, emergency and disaster management. Its origins begin with efforts from the Pan American Health Organization (PAHO) and the Latin America countries, but its reach has spread, and the HSI was applied extensively in other regions (including Europe) after the global campaign for hospitals safe from disasters [18,33,34]. Calculating the safety score allows the hospital to establish maintenance and monitoring routines and consider various necessary measures to improve safety in the medium term.

The total score for the healthcare facility can be in one of the three classifications regarding safety:

- ***Classification A***: Considered to be able to safely continue their activities in case of disasters (safety index 0.66-1.00)
- ***Classification B***: Considered to be able to resist against a crisis, but their equipment and vital services are exposed to danger (safety index 0.36-0.65)
- ***Classification C***: Considered to be unsafe for people working there and patients in crisis, requiring urgent intervention measures (safety index 0.00-0.35)

The information necessary for the calculation of the HSI and its evaluation in the context of a primary healthcare centre was collected through several structured interviews, conducted in 2017 with the Director of the PHC Obrenovac, the Technical Director, Chief of Occupational Safety and Health Department, and the President of the Emergency Response Team of the Obrenovac municipality. An unofficial translation of the HSI into the Serbian language was used, as no official translation exists.

### 2.3. Calculation

The total score and scores for each module were calculated using the Hospital Safety Index excel file (July 2017 version), which was kindly provided to us by the WHO Health Emergencies Programme office (Geneva, Switzerland, personal communication). The Excel file contains four sheets. The first one contains the weighted contribution of each question to the corresponding module, the second sheet is the questionnaire, the third sheet shows the summary of safety ratings, and the fourth sheet contains the inform on module-specific safety index and the overall hospital safety index. Each module’s contribution to the overall safety index was set to 1/3 (equal contribution of all three modules).

### 2.4. Evaluation of the Hospital safety index as a tool for primary healthcare centers’ safety assessment

During the structured interviews with the PHC Obrenovac management and staff, each of the HSI modules, as well as all underlying questions were evaluated as for their relevance for the functioning of a primary healthcare facility. Three levels of relevance were attributed to each question:

- ***High***: questions which have the same relevance for primary healthcare centers as for hospitals during disasters
- ***Low***: questions which have lower relevance for primary healthcare centers than for hospitals during disasters
- ***Not relevant (NR)***: questions which are not relevant in primary healthcare settings.

## 3. Results

Being a multi-risk assessment tool, the first part of the HSI evaluates the hazards, which could affect the safety of the hospital and the role of the hospital in emergency and disaster management. Floods have been identified as the main hazard that could affect the hospital. Earthquakes and landslides could also be of interest and have a history of occurring in Serbia. Other geological, hydro-meteorological, biological, technological, and societal hazards were not considered of importance for this PHC. Each of the HSI questions was rated by the study team according to their relevance to the primary healthcare center’s functioning during a disaster. Three levels of relevance: “High,” “Low,” and “Not relevant” were attributed to each question (see ***Section 2.4***.).

### 3.1. Overall hospital safety

***Table 1*** shows the overall safety index, the indices for structural safety, nonstructural safety, and emergency and disaster management, together with the number of items falling into the three safety categories (see ***Section 2.2***.). The overall safety of PHC Obrenovac fell into the A category, with an overall safety index of 0.82. Structural safety, nonstructural safety, and emergency and disaster management achieved scores of 0.95, 0.74, and 0.75, respectively.

**Table 1.** Hospital Safety Index evaluation of PHC Obrenovac

| Index | Number of items <sup>a</sup> |  |  | Crude Safety Index <sup>b</sup> | Safety Index <sup>c</sup> | Category |
| --- | --- | --- | --- | --- | --- | --- |
|  | Low | Average | High |  |  |  |
| **Overall Safety Index** | **17** | **24** | **107** | **/** | **0.82** | **A** |
| Structural safety | 0 | 1 | 17 | 0.96 | 0.95 | A |
| Nonstructural safety | 11 | 14 | 65 | 0.81 | 0.74 | A |
| Emergency and disaster management | 6 | 9 | 25 | 0.81 | 0.75 | A |

<sup>a</sup> Low = Unlikely to function, Average = Likely to function, High = Highly likely to function
<sup>b</sup> Non-bias adjusted safety index
<sup>c</sup> Bias-free safety index

### 3.2. Structural safety

This module has been designed to be used to address the structural elements, such as columns, beams, walls, floor slabs, foundations that form part of the load-bearing system of the buildings. Good structural safety implies that there is a low probability any of the above-mentioned elements would fail in case of a disaster.

***Table 2*** shows the overview of the structural safety HSI results of PHC Obrenovac. The structural safety of this health facility achieved the highest score among the modules (0.95) with no items with low safety, only one item with average safety and 17 items with high safety. Only structural system design (under the building integrity category) was scored as “average.”

**Table 2.** Structural safety of the PHC

| Items | Number of items |  |  |
| --- | --- | --- | --- |
|  | Low | Average | High |
| Prior events and hazards affecting building safety | 0 | 0 | 3 |
| Building integrity | 0 | 1 | 14 |
| **Total** | **0** | **1** | **17** |

All questions in the structural safety module were considered of “High” relevance. As this module evaluates the possibility of structural failure (e.g., collapse of the building) during a disaster, there were no differences found in the relevance of individual questions between hospitals and primary healthcare centers.

### 3.3. Nonstructural safety

The nonstructural safety module of the HSI includes four submodules: architectural safety; infrastructure protection, access, and physical security; critical systems; and equipment and supplies. These elements are considered critical to the functioning of the hospital, but do not belong to the structural component of the HSI, which is concentrated more on the load-bearing system of the hospital buildings.

***Table 3*** shows the overview of the nonstructural safety HSI results of PHC Obrenovac. The majority of items in all sub-categories were scored as high, although several sub-categories had up to 15 items evaluated as low or average, mostly in the architectural safety and critical systems categories. In architectural safety, four items were scored as average, while in critical systems, 8 and 7 items were evaluated as average and low, respectively. The “unlikely to function” items were in the categories of electrical, telecommunications, water supply systems. In the equipment and supplies, category 2 and 4 items were evaluated as average and low, respectively, located in the categories of office and storeroom furnishings and equipment. Insecure furnishings and equipment in offices and storerooms, which could pose a hazard in case of disasters, as well as the lack of medical and laboratory equipment and supplies used for diagnosis and treatment of patients to guarantee at least 72 hours of uninterrupted service at the maximum capacity, were also identified as weaknesses of PHC Obrenovac.

**Table 3.** Nonstructural safety of the PHC

| Items | Number of items |  |  |
| --- | --- | --- | --- |
|  | Low | Average | High |
| Architectural safety | 0 | 4 | 9 |
| Infrastructure protection, access, and physical security | 0 | 0 | 4 |
| Critical systems (subtotal) | 7 | 8 | 37 |
| Electrical systems | 1 | 1 | 8 |
| Telecommunication systems | 2 | 2 | 4 |
| Water supply system | 3 | 0 | 2 |
| Fire protection system | 0 | 2 | 3 |
| Waste management systems | 0 | 0 | 5 |
| Fuel storage systems | 0 | 0 | 5 |
| Medical gases systems | 0 | 1 | 5 |
| Heating, ventilation, and air-conditioning (HVAC) systems | 1 | 2 | 5 |
| Equipment and supplies (subtotal) | 4 | 2 | 15 |
| Office and storeroom furnishings and equipment (fixed and movable) | 2 | 0 | 0 |
| Medical and laboratory equipment and supplies used for diagnosis and treatment | 2 | 2 | 15 |
| **Total** | **11** | **14** | **65** |

The majority of items in the module on nonstructural safety were considered of the same relevance for primary healthcare centers as they are for hospitals. Differences were seen in the “medical gases system” category where all questions were marked as of “low” relevance. There were notable differences in the “medical and laboratory equipment and supplies used for diagnosis and treatment” category, where most of the questions were rated as “low” or “not relevant” for primary healthcare center’s functioning.

### 3.4. Emergency and disaster management

***Table 4*** shows the overview of the emergency and disaster management HSI results of PHC Obrenovac. The majority of items in various categories were scored as “highly likely to function,” although several categories contained items which point to improvements to be made. All items in the coordination of emergency and disaster management activities category have been scored as “high,” which underlines the PHC’s readiness for disasters from the management point of view. Most items not likely to function were found in the evacuation, decontamination and security category, while most items scored as average were found in the hospital emergency and disaster response planning, logistics and finance, and human resources categories.

**Table 4.** Emergency and disaster management of the PHC

| Items | Number of items |  |  |
| --- | --- | --- | --- |
|  | Low | Average | High |
| Coordination of emergency and disaster management activities | 0 | 0 | 8 |
| Hospital emergency and disaster response planning | 1 | 3 | 1 |
| Communication and information management | 0 | 0 | 4 |
| Human resources | 0 | 2 | 3 |
| Logistics and finance | 1 | 3 | 0 |
| Patient care and support services | 1 | 1 | 7 |
| Evacuation, decontamination and security | 3 | 0 | 2 |
| **Total** | **6** | **9** | **25** |

In the “emergency and disaster management” module, four items were marked as of “low” relevance in the “patient care and support services” and “evacuation, decontamination, and security” categories (2 in each category).

The evaluation of each question regarding the relevance to primary health care services is presented in ***Supplementary Tables S1, S2***, and ***S3***.

## 4. Discussion

In May 2014, floods struck Serbia, with the municipality of Obrenovac most severely affected. By the end of the day one, healthcare services provided by PHC Obrenovac stopped completely. Water barriers disabled transport around, in and out of the city, with an increased need for primary healthcare services, among which the highest demand came from chronic disease patients, people with diabetes, especially hemodialysis patients, and injured people. This study analyzes the safety and disaster preparedness of the PHC in Obrenovac using the Hospital Safety Index, a tool developed by the WHO to rapidly assess the safety of hospitals.

The structural, nonstructural, and emergency and disaster management modules were scored relatively high, putting PHC Obrenovac into the “A” category for safety. Regardless of the overall score, much was learned taking a more detailed look at the individual modules, with their categories, sub-categories, and questions. The building of PHC Obrenovac was constructed in 1952 following the structural safety standards of Yugoslavia at that time. Nevertheless, its structural safety received the highest score of 0.95. There was no significant structural damage to the building due to the floods in the past, and structural safety was not considered a vulnerability of the PHC. ***Figure 2*** shows the flooded PHC building during the floods of 2014, and although the building was flooded, no structural safety problems were identified. In a case study from Sudan, structural safety, reflected in significant damage by flooding and need for renovation, contributed to one-third of the vulnerability of the buildings [25]. Investments in hospitals’ physical safety, in a study from Iran, resulted in significant improvements in this area in just three years (2012 to 2015), moving the average safety score from 34 to 43 [14]. In fact, what the use of the HSI in this setting revealed was that the failure of the PHC to function during the 2014 floods was due to functional failures, which are considered a cornerstone of a health facility’s preparedness [35].

**Figure 2.**
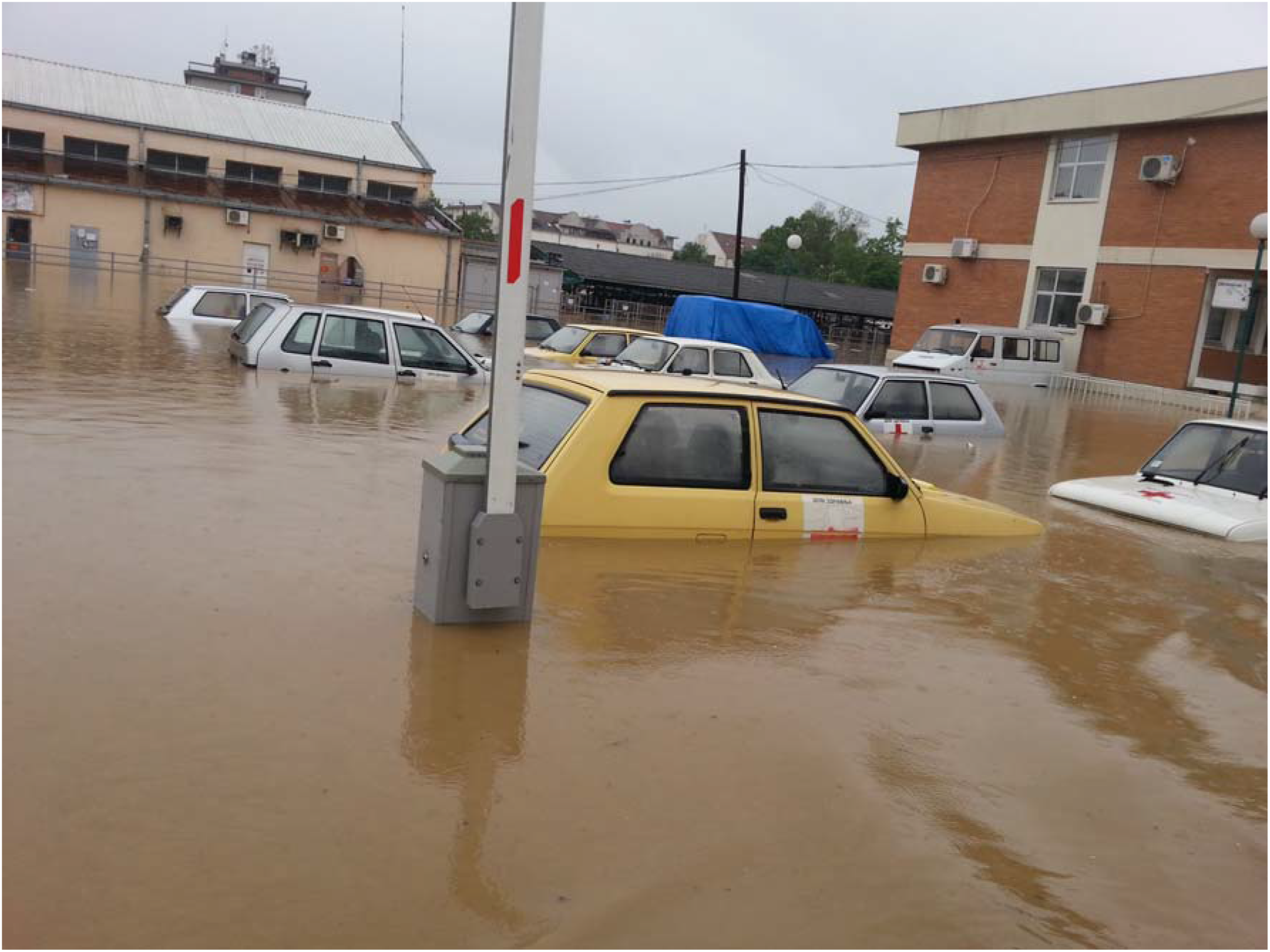
The flooded parking lot and Primary Healthcare Centre’s building during the floods of 2014.

Primary healthcare center Obrenovac received a score of 0.74 in the nonstructural safety module of the HSI, which is considered a high score, although in this module a number of items were scored “low” or “average.” In architectural safety, some items were scored as average due to the damage the building had sustained during the 2014 floods, and due to the fact it was built more than 60 years ago with irregular investments in repairs and improvements of the doors (above all exits and entrances), windows, shutters, and the roof. In critical systems, the safety score was not high due to the lack of resources to maintain and restore the electric power supply, and due to the fact, there is only one entrance for the local power supply.

Safety was estimated as low for the condition of the telecommunication systems and lack of alternative communications systems, and due to inadequate condition and protection of the external and internal communications systems. Most low ratings were given to the water supply system, as the water reserves would not allow 72 hours of functioning, the supplementary pumping system lacks, and problems are expected in the restoration of water supply. This analysis has underlined the importance of an adequate water supply system, as it is considered one of the main factors impacting the provision of health services [36,37]. In a study of water and power supply in the Greek islands, only half had water backup systems that would last for at least 72 hours (mostly in hospitals), with lower level health facilities less likely to have emergency water supplies [38].

In critical systems, the HSI has identified the backup power supply and its location as a weak point of PHC Obrenovac. In fact, during the 2014 floods, the backup power supply was rendered useless early, as it was located at the lowest level of the building, which was flooded first. The same lesson was learned from the 2002 Dresden flooding in Germany, where the authors noted that the power supply should not be positioned in places that are prone to flooding [39].

Another weak point, telecommunications, was also obvious during the floods, when all landlines, mobile phones, and the radio connection were lost as the electricity was out in the whole city, and the medical staff, together with the evacuated people, had no communication with the outside world except through rescue boats evacuating people from their homes. In a wide evaluation of the safety of hospitals in Iran, all health centers were found to have a fully functional and ready communication system in case of disasters [25]. This was achieved by a public-private partnership of the Ministry of Health and private communication companies, which provided a free of charge communication system to all health facilities down to the level of health centers. In the 21^st^ century, there is a vast quantity of technologies which can be used in flood disaster risk reduction, including social media, mapping platforms, and crowdsourcing, as well as many tools for live data collection, analysis and risk assessment [40].

It is well established that the availability of medical equipment and supplies is crucial for a hospital’s capacity during disasters [41–43], although data for PHCs is not available and could represent an interesting area of research [5]. During the 2014 floods in, all of the basement and ground floor rooms and services of PHC Obrenovac have been fully flooded by the end of day one, together with the storage of sanitary material and pharmaceutical products, as well as the cleaning and technical services (see ***Figures 3*** and ***4***). In a semi-quantitative risk assessment model of primary health care service interruption during floods, the availability of essential drugs and supplies has been underlined as one of the important variables of a facility’s capacity to cope with a disaster [25]. It is interesting to note that, in a study utilizing HSI, the lowest safety index for hospitals in Teheran (Iran) was found in the medical and laboratory equipment and supplies used for diagnosis and treatment, compared to the lowest safety index in the critical systems in Stockholm (Sweden) hospitals [15].

**Figure 3.**
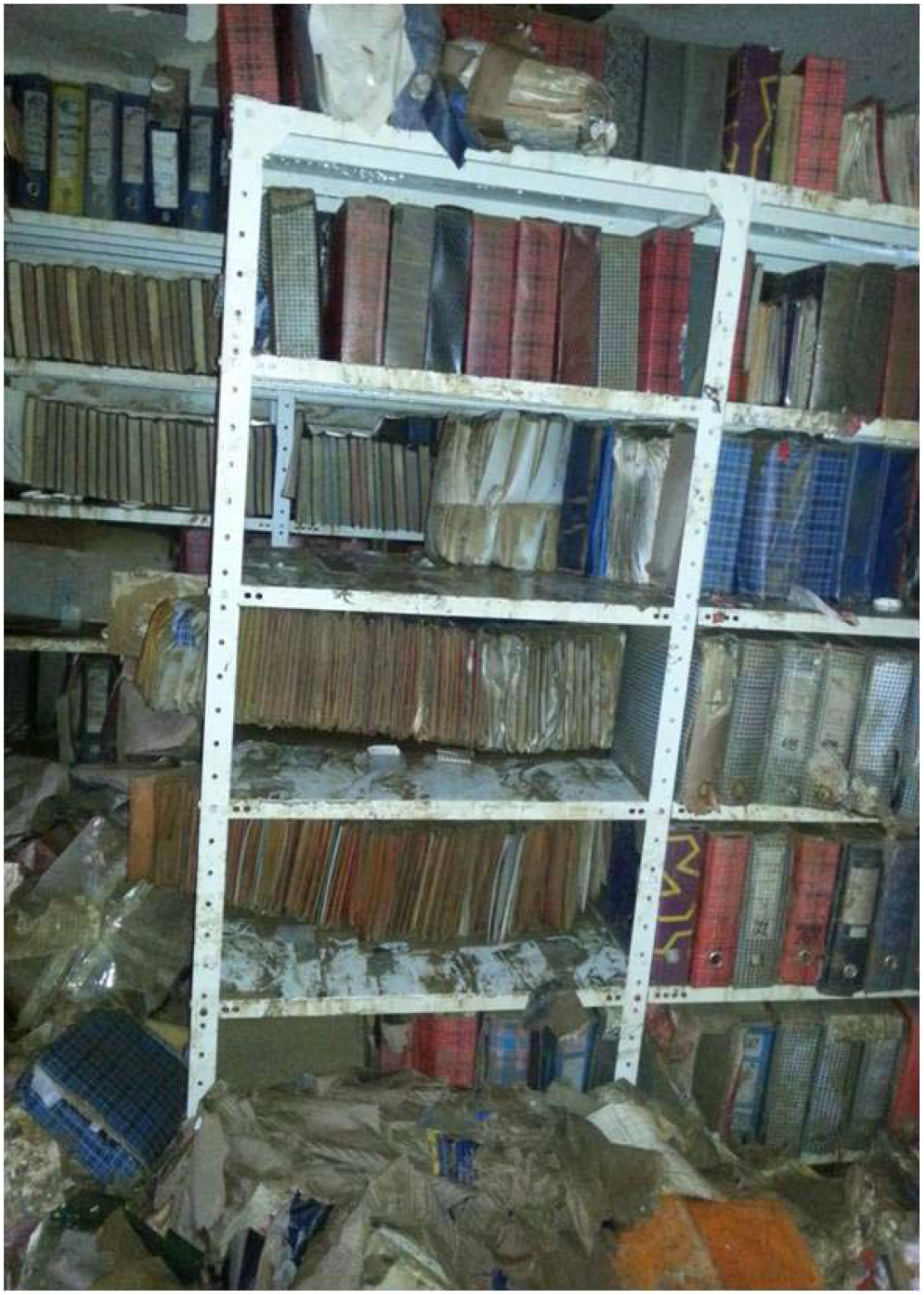
Archive of the Primary Healthcare Centre.

**Figure 4.**
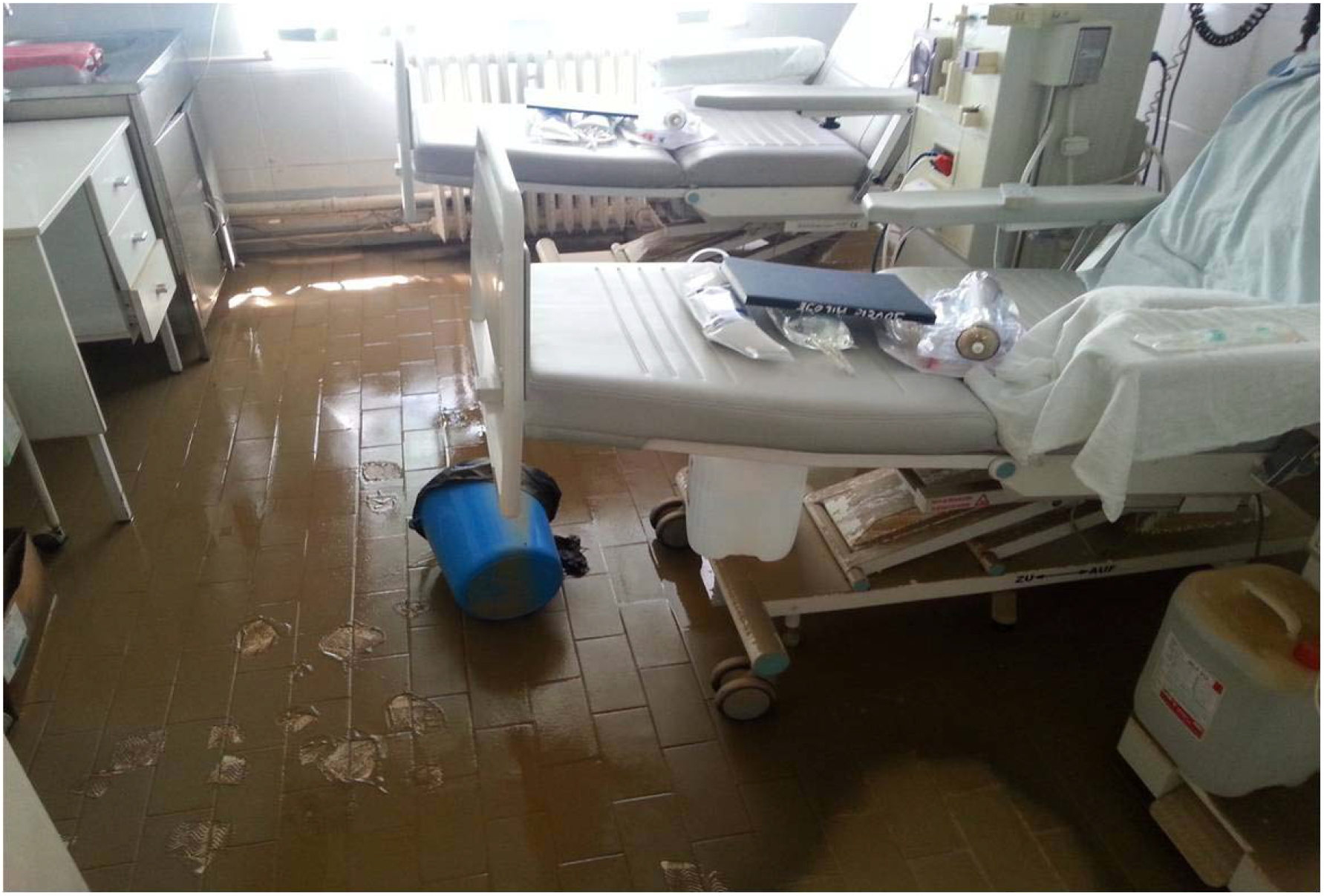
Hemodialysis room of the Primary Healthcare Centre.

Emergency and disaster management module of the HSI has demonstrated the adequate preparedness of PHC Obrenovac for hazards, which could be attributed to the recent flood of 2014 and the activities done during and after this disaster. Nevertheless, improvements are still to be made to the hazard-specific sub plans, procedures to activate and deactivate these plans, emergency and disaster response plan exercises, as well as the hospital recovery plan.

Logistics and finance category might require most improvement, as arrangements with local suppliers and vendors, transportation, food and drinking water, and financial resources exist but are not considered operational by the emergency management team. During the 2014 disaster, 50-80% of staff has been available for work. Still, no guarantee exists that they would have space and wellbeing measures available for more than 72 hours of functioning. Only staff members who had to take care of family members (due to kindergartens and schools being closed) did not show up for work in May 2014. In a detailed survey of hospital employees’ attitudes and needs regarding work commitments during disasters, most (81%) were willing to respond in case of floods but underlined the importance of child and pet care, as well as phone and email access [44]. The importance of communication has been underlined before, but having in mind the need for child and pet care for hospital staff should be taken into account in DRR for primary healthcare centers. In addition, much improvement is needed in the field of personal protective equipment for the hospital staff in the case of chemical/biological hazards and isolation in case of epidemics.

There is no internationally recognized and validated method to evaluate the safety of primary healthcare centers in case of disasters, although the interruption of services offered by these facilities is considered an evenly important problem as emergency response in large hospitals [19]. The HSI covers most of the factors important for this kind of evaluation, but a detailed analysis of questions and their relevance for the primary care setting is lacking. Our study underlines the usefulness of the structural safety module in primary healthcare settings but demonstrates the need for further evaluation of the nonstructural safety and emergency and disaster management modules. Most differences in the relevance of various categories and questions between primary healthcare centers and hospitals were seen in the medical gases system, as well as in the medical and laboratory equipment and supplies used for diagnosis and treatment. This result is to be expected, as primary healthcare centers serve a different purpose than hospitals. In Serbia, health services are organized through primary, secondary and tertiary care, with only some overlap between the services provided by primary (PHCs) and secondary (Hospitals) service. A PHCs role in this system, with or without disasters, remains in the diagnostic, follow-up, and non-invasive treatment, as well as triage of patients requiring treatment in a secondary or tertiary healthcare facility. Having in mind these differences, it is our belief that a modified HSI could prove to be a valuable, validated tool for safety assessment of primary healthcare facilities.

Little or no work has been done for the evaluation of hospitals’ and primary healthcare facilities’ safety in Serbia, as well as the South East Europe region, although the need is evident [34]. The main weaknesses of the present work, the fact that only one PHC was evaluated and that HSI, a tool intended for hospital evaluation was used, have also resulted in important insights on the use of HSI for PHC evaluation and potential directions of its development. Experiences of PHCs during disasters, such as that of Obrenovac presented in this paper, can help in developing a methodology for the identification of a PHC’s role in disasters, as well as to better quantify the importance of various questions to the overall score [25]. Officially translating the HSI into different languages, organizing a self-assessment of hospitals and PHCs, similar to that done in Iran, and developing tools for easier scoring of the HSI could reveal the areas of safety where most work is needed [14]. Country- or region-specific hazard assessment (Module 1 of the HSI) could help understand hazards relevant for different areas of the country, and guide the evaluation of the safety of hospitals and PHCs. A detailed study, similar to the study of the electronic health records in post-Hurricane Sandy time, could shed much needed light on the population’s healthcare needs, as well as how and where those needs were met during the 2014 floods [45].

## 5. Conclusions

The 2014 flood’s total effect on health in Serbia was estimated at 5.7 million euros, with 3 million due to damages and 2.7 million due to losses, and the post-disaster needs for recovery and reconstruction were estimated to 7.1 million euros. In the 30 days from the beginning of the flood, the staff of PHC Obrenovac had done a total of 15,488 physical examinations, of which 1,697 in the emergency room, 1,549 in the pediatrics, 289 in the gynecological, 73 in the obstetrics, and around 11,000 in the internal medicine departments. More than 10,000 patients received pharmaceutical products from the humanitarian aid supplies, and almost 1,000 patients were vaccinated. This work, done in the most difficult of times, underlines the importance primary healthcare facilities for the communities to which they provide services.

In the current study, the Hospital Safety Index was, for the first time, used for the evaluation of a PHC. The results of PHC Obrenovac have revealed areas where improvements are most needed to secure adequate functioning of this facility during disasters, which has also underlined the usefulness of this instrument even in the primary healthcare setting. Items constituting HSI have been evaluated for their relevance in the safety evaluation of primary healthcare facilities, which could allow the development of a modified HSI, which could be used for this purpose, achieving the goal of safe primary healthcare facilities and no interruption of services in case of disasters.

## Acknowledgments

We acknowledge the kind help and information provided by the staff of the PHC Obrenovac, without which this research would have been impossible. In addition, we would like to acknowledge the help provided by Dr. Jostacio Lapitan of the Country Health Emergency Preparedness and International Health Regulations (CPI), World Health Organization Health Emergencies Programme.

Research presented in this paper has been conducted as part of the project TR 34009 financed by the Ministry of Education, Science and Technological Development of the Republic of Serbia.

---

**Supplementary Table S1.**
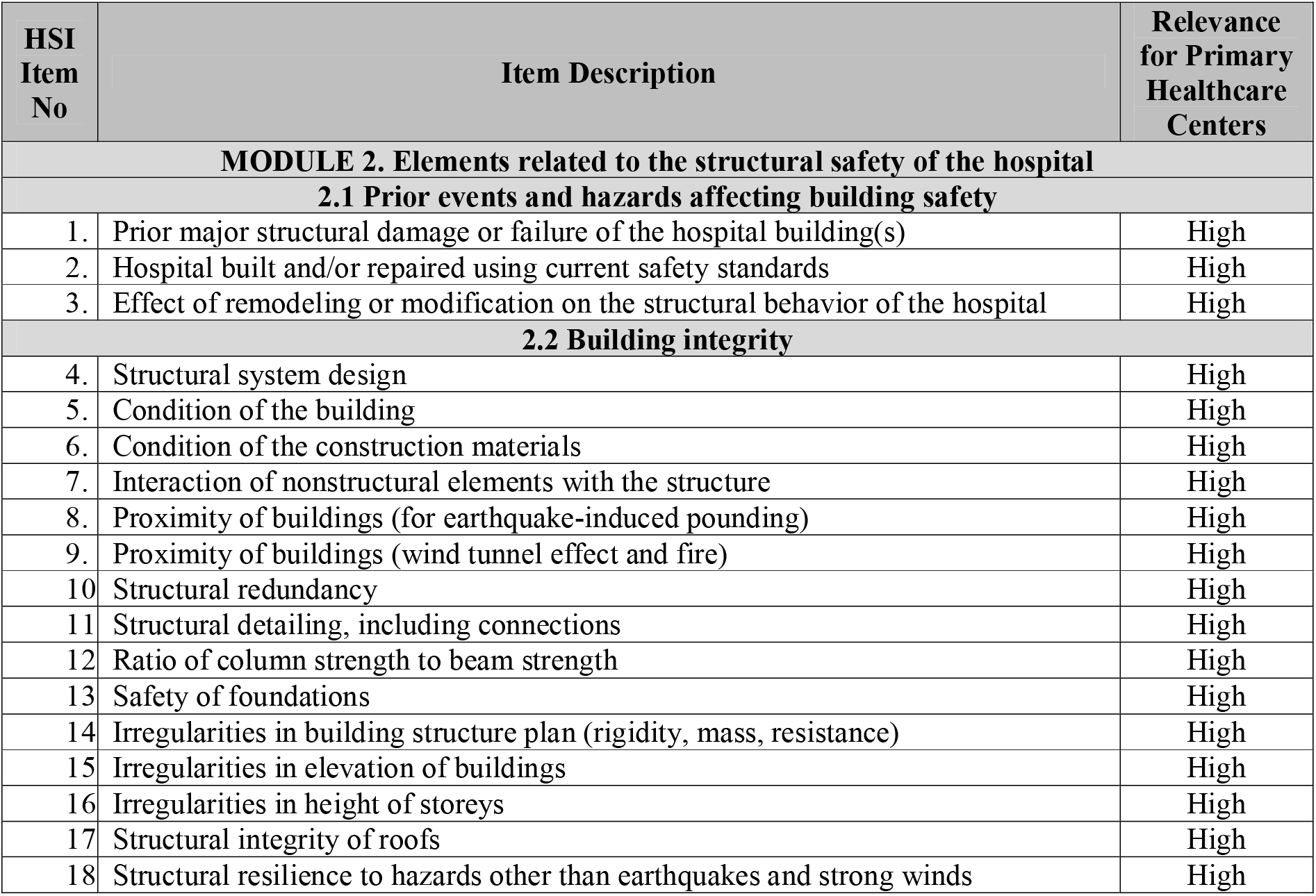
HSI Module 2 Items and their relevance in the primary healthcare setting.

**Supplementary Table S2.** HSI Module 3 Items and their relevance in the primary healthcare setting.

| HSI Item No | Item Description | Relevance for Primary Healthcare Centers |
| --- | --- | --- |
| **MODULE 3: Elements related to the non-structural safety of the hospital** |  |  |
| **3.1 Architectural safety** |  |  |
| 19. | Major damage and repair of the nonstructural elements | High |
| 20. | Condition and safety of doors, exits and entrances | High |
| 21. | Condition and safety of windows and shutters | High |
| 22. | Condition and safety of other elements of the building envelope (e.g. outside walls, facings) | High |
| 23. | Condition and safety of roofing | High |
| 24. | Condition and safety of railings and parapets | High |
| 25. | Condition and safety of perimeter walls and fencing | High |
| 26. | Condition and safety of other architectural elements (e.g. cornices, ornaments, chimneys, signs) | High |
| 27. | Safe conditions for movement outside the hospital buildings | High |
| 28. | Safe conditions for movement inside the building (e.g. corridors, stairs) | High |
| 29. | Condition and safety of internal walls and partitions | High |
| 30. | Condition and safety of false or suspended ceilings | High |
| 31. | Condition and safety of the elevator system | High |
| 32. | Condition and safety of stairways and ramps | High |
| 33. | Condition and safety of floor coverings | High |
| **3.2 Infrastructure protection, access and physical security** |  |  |
| 34. | Location of hospital's critical services and equipment in relation to local hazards | High |
| 35. | Hospital access routes | High |
| 36. | Emergency exits and evacuation routes | High |
| 37. | Physical security of building, equipment, staff and patients | High |
| **3.3 Critical systems** |  |  |
| **3.3.1 Electrical systems** |  |  |
| 38. | Capacity of alternative sources of electricity (e.g. generators) | High |
| 39. | Regular tests of alternative sources of electricity in critical areas | High |
| 40. | Condition and safety of alternative source(s) of electricity | High |
| 41. | Condition and safety of electrical equipment, cables and cable ducts | High |
| 42. | Redundant system for the local electric power supply | High |
| 43. | Condition and safety of control panels, overload breaker switches and cables | High |
| 44. | Lighting system for critical areas of the hospital | High |
| 45. | Condition and safety of internal and external lighting systems | High |
| 46. | External electrical systems installed for hospital usage | High |
| 47. | Emergency maintenance and restoration of electric power supply and alternative sources | High |
| **3.3.2 Telecommunication systems** |  |  |
| 48. | Condition and safety of antennas | High |
| 49. | Condition and safety of low- and extra-low-voltage systems (Internet and telephone) | High |
| 50. | Alternative communication systems | High |
| 51. | Condition and safety of telecommunications equipment and cables | High |
| 52. | Effect of external telecommunications systems on hospital communications | High |
| 53. | Safety of sites for telecommunication systems | High |
| 54. | Condition and safety of internal communications systems | High |
| 55. | Emergency maintenance and restoration of standard and alternate communications | High |
|  | systems |  |
| **3.3.3 Water supply system** |  |  |
| 56. | Water reserves for hospital services and functions | High |
| 57. | Location of water storage tanks | High |
| 58. | Safety of the water distribution system | High |
| 59. | Alternative water supply to the regular water supply | High |
| 60. | Supplementary pumping system | High |
| 61. | Emergency maintenance and restoration of water supply | High |
| **3.3.4 Fire protection system** |  |  |
| 62. | Condition and safety of the fire protection (passive) system | High |
| 63. | Fire/smoke detection systems | High |
| 64. | Fire suppression systems (automatic and manual) | High |
| 65. | Water supply for fire suppression | High |
| 66. | Emergency maintenance and restoration of the fire protection system |  |
| **3.3.5 Waste management systems** |  |  |
| 67. | Safety of nonhazardous wastewater systems | High |
| 68. | Safety of hazardous wastewater and liquid waste | High |
| 69. | Safety of nonhazardous solid waste system | High |
| 70. | Safety of hazardous solid waste system | High |
| 71. | Emergency maintenance and restoration of all types of hospital waste management systems | High |
| **3.3.6 Fuel storage systems (e.g. gas, gasoline and diesel)** |  |  |
| 72. | Fuel reserves | High |
| 73. | Condition and safety of above-ground fuel tanks and/or cylinders | High |
| 74. | Safe location of fuel storage away from hospital buildings | High |
| 75. | Condition and safety of the fuel distribution system (valves, hoses, and connections) | High |
| 76. | Emergency maintenance and restoration of fuel reserves | High |
| **3.3.7 Medical gases systems** |  |  |
| 77. | Location of storage areas for medical gases | Low |
| 78. | Safety of storage areas for the medical gas tanks and/or cylinders | Low |
| 79. | Condition and safety of medical gas distribution system (e.g. valves, pipes, connections) | Low |
| 80. | Condition and safety of medical gas cylinders and related equipment in the hospital | Low |
| 81. | Availability of alternative sources of medical gases | Low |
| 82. | Emergency maintenance and restoration of medical gas systems. | Low |
| **3.3.8 Heating, ventilation, and air-conditioning (HVAC) systems** |  |  |
| 83. | Adequate location of enclosures for HVAC equipment | High |
| 84. | Safety of enclosures for HVAC equipment | High |
| 85. | Safety and operating condition of HVAC equipment (e.g. boiler, exhaust) | High |
| 86. | Adequate supports for ducts and review of flexibility of ducts and piping that cross expansion joints | High |
| 87. | Condition and safety of pipes, connections and valves | High |
| 88. | Condition and safety of air-conditioning equipment | High |
| 89. | Operation of air-conditioning system (including negative pressure areas) | High |
| 90. | Emergency maintenance and restoration of the HVAC systems | High |
| **3.4 Equipment and supplies** |  |  |
| **3.4.1 Office and storeroom furnishings and equipment (fixed and movable)** |  |  |

| **HSI**<br>**Item**<br>**No** | **Item Description** | **Relevance**<br>**for Primary**<br>**Healthcare**<br>**Centers** |
| --- | --- | --- |
| 91. | Safety of shelving and shelf contents | High |
| 92. | Safety of computers and printers | High |
| **3.4.2 Medical and laboratory equipment and supplies used for diagnosis and treatment** |  |  |
| 93. | Safety of medical equipment in operating theatres and recovery rooms | Low |
| 94. | Condition and safety of radiology and imaging equipment | High |
| 95. | Condition and safety of laboratory equipment and supplies | High |
| 96. | Condition and safety of medical equipment in emergency care services unit | High |
| 97. | Condition and safety of medical equipment in intensive or intermediate care unit | NR |
| 98. | Condition and safety of equipment and furnishings in the pharmacy | High |
| 99. | Condition and safety of equipment and supplies in the sterilization services | High |
| 100. | Condition and safety of medical equipment for obstetric emergencies and neonatal care | NR – Low |
| 101. | Condition and safety of medical equipment and supplies emergency care for burns | Low |
| 102. | Condition and safety of medical equipment for nuclear medicine and radiation therapy | NR |
| 103. | Condition and safety of medical equipment in other services | High |
| 104. | Medicines and supplies | High |
| 105. | Sterilized instruments and other materials | High |
| 106. | Medical equipment specifically used in emergencies and disasters | High |
| 107. | Supply of medical gases | Low |
| 108. | Mechanical volume ventilators | NR |
| 109. | Electromedical equipment | NR |
| 110. | Life-support equipment | NR |
| 111. | Supplies, equipment or crash carts for cardiopulmonary arrest | Low |

**Supplementary Table S3.** HSI Module 4 Items and their relevance in the primary healthcare setting.

| No | HSI Item Description | Relevance for Primary Healthcare Centers |
| --- | --- | --- |
| **MODULE 4. Emergency and disaster management** |  |  |
| **4.1 Coordination of emergency and disaster management activities** |  |  |
| 112. | Hospital Emergency/Disaster Committee | High |
| 113. | Committee member responsibilities and training | High |
| 114. | Designated emergency and disaster management coordinator | High |
| 115. | Preparedness programme for strengthening emergency and disaster response and recovery | High |
| 116. | Hospital incident management system | High |
| 117. | Emergency Operations Centre (EOC) | High |
| 118. | Coordination mechanisms and cooperative arrangements with local emergency/disaster management agencies | High |
| 119. | Coordination mechanisms and cooperative arrangements with the health-care network | High |
| **4.2 Hospital emergency and disaster management response and recovery planning** |  |  |
| 120. | Hospital emergency or disaster response plan | High |
| 121. | Hazard-specific subplans | High |
| 122. | Procedures to activate and deactivate plans | High |
| 123. | Hospital emergency and disaster response plan exercises, evaluation and corrective actions | High |
| 124. | Hospital recovery plan |  |
| **4.3 Communication and information management** |  |  |
| 125. | Emergency internal and external communication | High |
| 126. | External stakeholder directory | High |
| 127. | Procedures for communicating with the public and media | High |
| 128. | Management of patient information | High |
| **4.4 Human resources** |  |  |
| 129. | Staff contact list | High |
| 130. | Staff availability | High |
| 131. | Mobilization and recruitment of personnel during an emergency or disaster | High |
| 132. | Duties assigned to personnel for emergency or disaster response and recovery | High |
| 133. | Well-being of hospital personnel during an emergency or disaster | High |
| **4.5 Logistics and finance** |  |  |
| 134. | Agreements with local suppliers and vendors for emergencies and disasters | High |
| 135. | Transportation during an emergency | High |
| 136. | Food and drinking-water during an emergency | High |
| 137. | Financial resources for emergencies and disasters | High |
| **4.6 Patient care and support services** |  |  |
| 138. | Continuity of emergency and critical care services | High |
| 139. | Continuity of essential clinical support services | High |
| 140. | Expansion of usable space for mass casualty incidents | High |
| 141. | Triage for major emergencies and disasters | High |
| 142. | Triage tags and other logistical supplies for mass casualty incidents | Low |
| 143. | System for referral, transfer and reception of patients | High |
| 144. | Infection surveillance, prevention, and control procedures | Low |
| 145. | Psychosocial services | High |
| 146. | Post-mortem procedures in a mass fatality incident | High |
| **4.7 Evacuation, decontamination and security** |  |  |

| No | HSI Item Description | Relevance<br>for Primary<br>Healthcare<br>Centers |
| --- | --- | --- |
| 147. | Evacuation plan | High |
| 148. | Decontamination for chemical/biological hazards | Low |
| 149. | Personal protection equipment and isolation for infectious diseases and epidemics | Low |
| 150. | Emergency security procedures | High |
| 151. | Computer system network security | Low |

## Notes

**Declarations of interest:** none

## References

[1] UNISDR, Hyogo Framework for Action 2005-2015, in: United Nations Int. Strateg. Disaster Reduc, 2005. doi:10.1017/CBO9781107415324.004.

[2] A. Aitsi-Selmi, S. Egawa, H. Sasaki, C. Wannous, V. Murray, The Sendai framework for disaster risk reduction: Renewing the global commitment to people’s resilience, health, and well-being, Int. J. Disaster Risk Sci. 6 (2015) 164–176.

[3] A.S. Elnashai, L. Di Sarno, Fundamentals of earthquake engineering, Wiley New York, 2008.

[4] K.I. Shoaf, S.J. Rottman, Public health impact of disasters, Aust. J. Emerg. Manag. 15 (2000) 58–63.

[5] T. Jakubicka, F. Vos, R. Phalkey, D. Guha-Sapir, M. Marx, Health impacts of floods in Europe: Data gaps and information needs from a spatial perspective, Centre for Research on the Epidemiology of Disasters (CRED), 2010.

[6] R.H. Ofrin, I. Nelwan, Disaster risk reduction through strengthened primary health care, in: Reg. Heal. Forum, World Health Organization Regional Office for South-East Asia, 2009: p. 29.

[7] W. Higgins, C. Wainright, N. Lu, R. Carrico, Assessing hospital preparedness using an instrument based on the Mass Casualty Disaster Plan Checklist: Results of a statewide survey, Am. J. Infect. Control. 32 (2004) 327–332. doi:10.1016/j.ajic.2004.03.006.

[8] B. Adini, A. Goldberg, D. Laor, R. Cohen, R. Zadok, Y. Bar-Dayan, Assessing levels of hospital emergency preparedness, Prehosp. Disaster Med. 21 (2006) 451–457. doi:10.1017/S1049023X00004192.

[9] A.H. Kaji, V. Langford, R.J. Lewis, Assessing Hospital Disaster Preparedness: A Comparison of an On-Site Survey, Directly Observed Drill Performance, and Video Analysis of Teamwork, Ann. Emerg. Med. 52 (2008). doi:10.1016/j.annemergmed.2007.10.026.

[10] E.J. Lazar, N. V. Cagliuso, K.M. Gebbie, Are we ready and how do we know? the urgent need for performance metrics in hospital emergency management, Disaster Med. Public Health Prep. 3 (2009) 57–60. doi:10.1097/DMP.0b013e31817e0e7f.

[11] M. Top, Ö. Gider, Y. Tas, An Investigation of Hospital Disaster Preparedness in Turkey, J. Homel. Secur. Emerg. Manag. 7 (2010). doi:10.2202/1547-7355.1781.

[12] W.H. Organization, Hospital safety index: Guide for evaluators, World Health Organization, 2015.

[13] A. Ardalan, Hospitals Safety from Disasters in I.R.Iran: The Results from Assessment of 224 Hospitals, PLOS Curr. (2014) 1–16. doi:10.1371/currents.dis.8297b528bd45975bc6291804747ee5db.

[14] A. Ardalan, M.K. Keleh, A. Saberinia, D. Khorasani-Zavareh, H. Khankeh, J. Miadfar, S. Maleknia, A. Mobini, S. Mehranamin, 2015 Estimation of Hospitals Safety from disasters in I.R.Iran: The results from the assessment of 421 hospitals, PLoS One. 11 (2016) 1–10. doi:10.1371/journal.pone.0161542.

[15] A. Djalali, A. Ardalan, G. Ohlen, P.L. Ingrassia, F. Della Corte, M. Castren, L. Kurland, Nonstructural safety of hospitals for disasters: A comparison between two capital cities, Disaster Med. Public Health Prep. 8 (2014) 179–184. doi:10.1017/dmp.2014.21.

[16] K. Jahangiri, Y.O. Izadkhah, A. Lari, Hospital safety index (HSI) analysis in confronting disasters: A case study from Iran, Int. J. Heal. Syst. Disaster Manag. 2 (2014) 44. doi:10.4103/2347-9019.135368.

[17] R. Miniati, C. Iasio, Vulnerability to Earthquakes and Floods of the Healthcare System in Florence, Italy, 2014. doi:10.1016/B978-0-12-410528-7.00004-7.

[18] A. Aiello, M. Pecce, L. Di Sarno, D. Perrone, F. Rossi, A safety index for hospital buildings, Disaster Adv. 5 (2012) 270–277.

[19] G.A. Mensah, A.H. Mokdad, S.F. Posner, E. Reed, E.J. Simoes, M.M. Engelgau, V.P. in N.D.W. Group, When chronic conditions become acute: prevention and control of chronic diseases and adverse health outcomes during natural disasters, Prev. Chronic Dis. 2 (2005).

[20] R.C. Kessler, P.S. Wang, D. Kendrick, N. Lurie, B. Springgate, Hurricane Katrina’s impact on the care of survivors with chronic medical conditions, J. Gen. Intern. Med. 22 (2007) 1225–1230. doi:10.1007/s11606-007-0294-1.

[21] K. Kamoi, M. Tanaka, T. Ikarashi, M. Miyakoshi, Effect of the 2004 Mid Niigata Prefecture earthquake on glycemic control in type 1 diabetic patients, Diabetes Res. Clin. Pract. 74 (2006) 141–147. doi:10.1016/j.diabres.2006.03.028.

[22] A. Inui, H. Kitaoka, M. Majima, S. Takamiya, M. Uemoto, C. Yonenaga, M. Honda, K. Shirakawa, N. Ueno, K. Amano, S. Morita, A. Kawara, K. Yokono, M. Kasuga, H. Taniguchi, Effect of the Kobe earthquake on stress and glycemic control in patients with diabetes mellitus, Arch. Intern. Med. 158 (1998) 274–278. doi:10.1001/archinte.158.3.274.

[23] K. Kario, T. Matsuo, H. Kobayashi, K. Yamamoto, K. Shimada, Earthquake-induced potentiation of acute risk factors in hypertensive elderly patients: Possible triggering of cardiovascular events after a major earthquake, J. Am. Coll. Cardiol. 29 (1997) 926–933. doi:10.1016/S0735-1097(97)00002-8.

[24] L. Redwood-Campbell, J. Abrahams, Primary Health care and Disasters—The Current State of the Literature: What We Know, Gaps and Next Steps, Prehosp. Disaster Med. 26 (2011) 184–191. doi:10.1017/S1049023X11006388.

[25] H.B. Abbas, J.K. Routray, A semi-quantitative risk assessment model of primary health care service interruption during flood: Case study of Aroma locality, Kassala State of Sudan, Int. J. Disaster Risk Reduct. 6 (2013) 118–128. doi:10.1016/j.ijdrr.2013.10.002.

[26] W. Du, G.J. Fitzgerald, M. Clark, X.Y. Hou, Health impacts of floods, Prehosp. Disaster Med. 25 (2010) 265–272. doi:10.1017/S1049023X00008141.

[27] M. Ahern, R.S. Kovats, P. Wilkinson, R. Few, F. Matthies, Global health impacts of floods: epidemiologic evidence, Epidemiol. Rev. 27 (2005) 36. doi:10.1093/epirev/mxi004.

[28] M. Wahlstrom, D. Guha-Sapir, The human cost of weather-related disasters 1995–2015, Geneva United Nations Int. Strateg. Disaster Reduct. (2015).

[29] G. Greenough, M. McGeehin, S.M. Bernard, J. Trtanj, J. Riad, D. Engelberg, The potential impacts of climate variability and change on health impacts of extreme weather events in the United States, Env. Heal. Perspect. 109 Suppl (2001) 191–198. doi:10.2307/3435009.

[30] T. Madsen, E. Figdor, When it rains, it pours: global warming and the rising frequency of extreme precipitation in the United States, Environment Texas Research & Policy Center, 2007.

[31] S. Hajat, K.L. Ebi, R.S. Kovats, B. Menne, S. Edwards, A. Haines, The human health consequences of flooding in Europe: A review, in: Extrem. Weather Events Public Heal. Responses, 2005: pp. 185–196. doi:10.1007/3-540-28862-7_18.

[32] Serbian Environmental Protection Agency, Serbia Floods 2014, Belgrade, 2014. http://www.sepa.gov.rs/download/SerbiaRNAreport_2014.pdf.

[33] G. Rockenschaub, K. V Harbou, Disaster resilient hospitals: An essential for all-hazards emergency preparedness, World Hosp. Heal. Serv. J. Int. Hosp. Fed. Pap. from 38th IHF World Hosp. Congr. Oslo. 49 (2013) 28–30.

[34] V. Radovic, K. Vitale, P.B. Tchounwou, Health facilities safety in natural disasters: Experiences and challenges from south east Europe, Int. J. Environ. Res. Public Health. 9 (2012) 1677–1686. doi:10.3390/ijerph9051677.

[35] A. Djalali, M. Castren, H. Khankeh, D. Gryth, M. Radestad, G. Ohlen, L. Kurland, Hospital disaster preparedness as measured by functional capacity: a comparison between Iran and Sweden., Prehosp. Disaster Med. 28 (2013) 454–461. doi:10.1017/S1049023X13008807.

[36] R. Aghababian, C.P. Lewis, L. Gans, F.J. Curley, Disasters Within Hospitals, Ann. Emerg. Med. 23 (1994) 771–777. doi:10.1016/S0196-0644(94)70313-2.

[37] K. Tanaka, The Kobe earthquake: the system response. A disaster report from Japan, Eur. J. Emerg. Med. 3 (1996) 263–269. http://www.biomednet.com/db/medline/97209198.

[38] L.-C.G. Alexakis, T.A. Codreanu, S.J. Stratton, Water and Power Reserve Capacity of Health Facilities in the Greek Islands, Prehosp. Disaster Med. 29 (2014) 146–150. doi:10.1017/S1049023X14000077.

[39] D. Meusel, W. Kirch, Lessons to be learned from the 2002 floods in Dresden, Germany, in: Extrem. Weather Events Public Heal. Responses, 2005. doi:10.1007/3-540-28862-7_17.

[40] I. McCallum, W. Liu, L. See, R. Mechler, A. Keating, S. Hochrainer-Stigler, J. Mochizuki, S. Fritz, S. Dugar, M. Arestegui, M. Szoenyi, J.C.L. Bayas, P. Burek, A. French, I. Moorthy, Technologies to Support Community Flood Disaster Risk Reduction, Int. J. Disaster Risk Sci. 7 (2016) 198–204. doi:10.1007/s13753-016-0086-5.

[41] A. Kaji, K.L. Koenig, T. Bey, Surge Capacity for Healthcare Systems: A Conceptual Framework, Acad. Emerg. Med. 13 (2006) 1157–1159. doi:10.1197/j.aem.2006.06.032.

[42] D.F. Barbisch, K.L. Koenig, Understanding Surge Capacity: Essential Elements, Acad. Emerg. Med. 13 (2006) 1098–1102. doi:10.1197/j.aem.2006.06.041.

[43] J.L. Hick, K.L. Koenig, D. Barbisch, A.B. Tareg, Surge capacity concepts for health care facilities: The CO-S-TR model for initial incident assessment, Disaster Med. Public Health Prep. 2 (2008). doi:10.1097/DMP.0b013e31817fffe8.

[44] D.C. Cone, B. a Cummings, Hospital disaster staffing: if you call, will they come?, Am. J. Disaster Med. 1 (2006) 28–36. http://www.ncbi.nlm.nih.gov/pubmed/18274041.

[45] K. Sebek, L. Jacobson, J. Wang, R. Newton-Dame, J. Singer, Assessing capacity and disease burden in a virtual network of New York city primary care providers following hurricane sandy, J. Urban Heal. 91 (2014) 615–622. doi:10.1007/s11524-014-9874-7.

